# Integrative identification by Hi-C revealed distinct advanced structural variations in Lung Adenocarcinoma tissue

**DOI:** 10.1101/2020.10.04.325738

**Authors:** Li Zhang, Tingting Song, Menglin Yao, Ying Yang, Zhiqiang Liu, Weimin Li

## Abstract

Advanced three-dimensional structure variations of chromatin in large genome fragments, such as conversion of A/B compartment, topologically associated domains (TADs) and chromatin loops are related closely to occurrence of malignant tumors. However, the structural characteristics of lung cancer still remain uncovered. In this study, we used high-throughput chromosome (HiC) confirmation capture to detect 2 non-smoking female lung adenocarcinoma (LUAD) tumor and paired normal tissues. The results indicated that significant chromatin variations were detected in tumor tissues compared with normal tissues. At compartment scale, the main conversion type of compartment is A→B in tumor tissues, which concentrated mainly on chromosome 3 (33.6%). Total of 216 tumor-specific TADs were identified in tumor tissues, which distributed mainly in chr1 (19), chr2 (15) and chr3(17). Finally we observed 41 distinct enhancer-promoter loops in tumor tissue, which associated closely to tumor-related pathways including MAPK, PI3K-AKT, Ras, Wnt and Ras1. The most important observation in this study was that we identified 5 important genes (*SYT16, NCEH1, NXPE3, MB21D2*, and *DZIP1L*), which were detected in both A→B compartment, TADs and chromatin loops. Four of these genes (*NCEH1, NXPE3, MB21D2*, and *DZIP1L*) located on q arm of chromosome 3, which also revealed the importance of chr3 in occurrence of structural variations in LUAD.

## Introduction

Lung cancer is leading cause of malignant tumor related morbidity and mortality, non-small cell lung cancer (NSCLC) accounts for 80% in all lung cancer patients. Smoke is considered an important factor in inducing lung cancer, but 50% of women lung cancer patients are not to tobacco use. The occurrence of lung cancer involves multiple mutations of tumor-related genes, including TP53, EGFR, KRAS et al., which can dramatically promote tumor development. Besides mutations in onco- and onco-suppressive genes, structure variations caused by chromatin instability, such as deletion, reversion, translocation, insertion and fusion, also act as critical roles in lung cancer development (Edwards PA 2010). In recent years, advanced conformational variations in large genome fragments, including A/B compartment, topologically associating domains (TADs) and chromatin loops, are also considered important in tumoregenesis.

High-throughput chromosome (Hi-C) conformation capture is a novel technology combined with high-throughput sequencing and chromatin conformation capture technology for quantifying long-range physical interactions in the genome, which is widely used to detect large fragments and advanced conformational variations, including A/B compartment, TADs and chromatin loops (Spielmann M et al. 2018), which are the main types of advanced chromatin structure variation in cells. A/B compartment is that each chromosome occupies a separate space (chromosome territory) in the interphase nucleus and is partitioned into distinct neighborhoods or compartments (Lieberman-Aiden E et al. 2009). When analyzed at sufficient resolution (typically below 100 kb), Hi-C experiments reveal the existence of distinct structural domains, referred to as Topologically Associating Domains (TADs), are characterized by increased intradomain interactions and insulation from their surroundings (Barutcu AR et al. 2018; Chang LH et al. 2020). The chromatin loop is formed by chromatin interactions between distal cis-regulatory elements to allow modulation of promoter activity by enhancers (Grubert F et al. 2020; Rao SS et al. 2014).

HiC is now applied in multiple tumors to detect specific advanced chromatin variations. In multiple myeloma (MM) cell lines (RPMI-8226 andU266), combined with Chip-seq and whole genome sequence (WGS), more than 100 chromosome interactions are detected and the A/B compartment switch region related closely to MAPK signaling pathway and cytokine-cytokine receptor interaction pathway (Wu P et al. 2017). In prostate cancer, Hi-C and WGS are performed in normal prostate epithelial cells PrEC and two prostate cancer cell lines (PC3 and LNCaP), which revealed that TADs are more numerous in tumor cells and copy number variations associate with formation of new TADs (Taberlay PC et al. 2016). In another study about prostate cancer, it is also identified specific enhancer-promoter loops in prostate cancer cell line (C42B and 22Rv1) (Rhie SK et al. 2019). All these results suggest that advanced structure variations (A/B compartment, TADs and chromatin loop) may play important roles in tumor development.

However, our knowledge of advanced structure variations in malignant tumors is still scarce. Variation of chromatin interaction patterns among diverse tissue types remains poorly defined, and its functional relationship with gene regulation remains to be characterized. Previous studies on structure variations are almost tumor cell lines, not tumor tissues from clinical patients, while the molecular events in tumor cell lines can not reflect the real situation in tumor development.

In this study, we performed Hi-C sequence on tumor and adjacent normal tissues from 2 paired non-smoke female lung adnocarcinoma (LUAD) patients. Further analysis was performed to analyze the difference of A/B compartment, TADs and chromatin loops between tumor and normal tissues. We identified cancer-specific A/B compartment, TADs and loops in tumor tissues. Our results suggested that genomic structural variation of LUAD was mainly derived from chromosome 3 and five genes, *SYT16, NCEH1, NXPE3, MB21D2, DZIP1L*, were identified, which might act as critical functions in development of LUAD.

## Result

### HIC detection revealed multiple structural variations in non-smoke LUAD tumor tissues

Genome disorder contributed to occurrence and development of malignant tumors (Flavahan WA et al. 2016; Hnisz D et al. 2016). In this study we conducted HIC analysis on 2 non-smoke female LUAD tumor and adjacent normal tissues, their ages was 37 years old and the clinical pathological stages all were III stage (Fig. 1A). Hi-C was performed to sequence all two samples and ∼54 and ∼52 million interactions were obtained from the cancer and normal tissues after filtering by_the ratio of invalid interaction pairs, which was analyzed by HiC-Pro software based on unique reads mapped on genome (Table S1).

**Figure 1.**
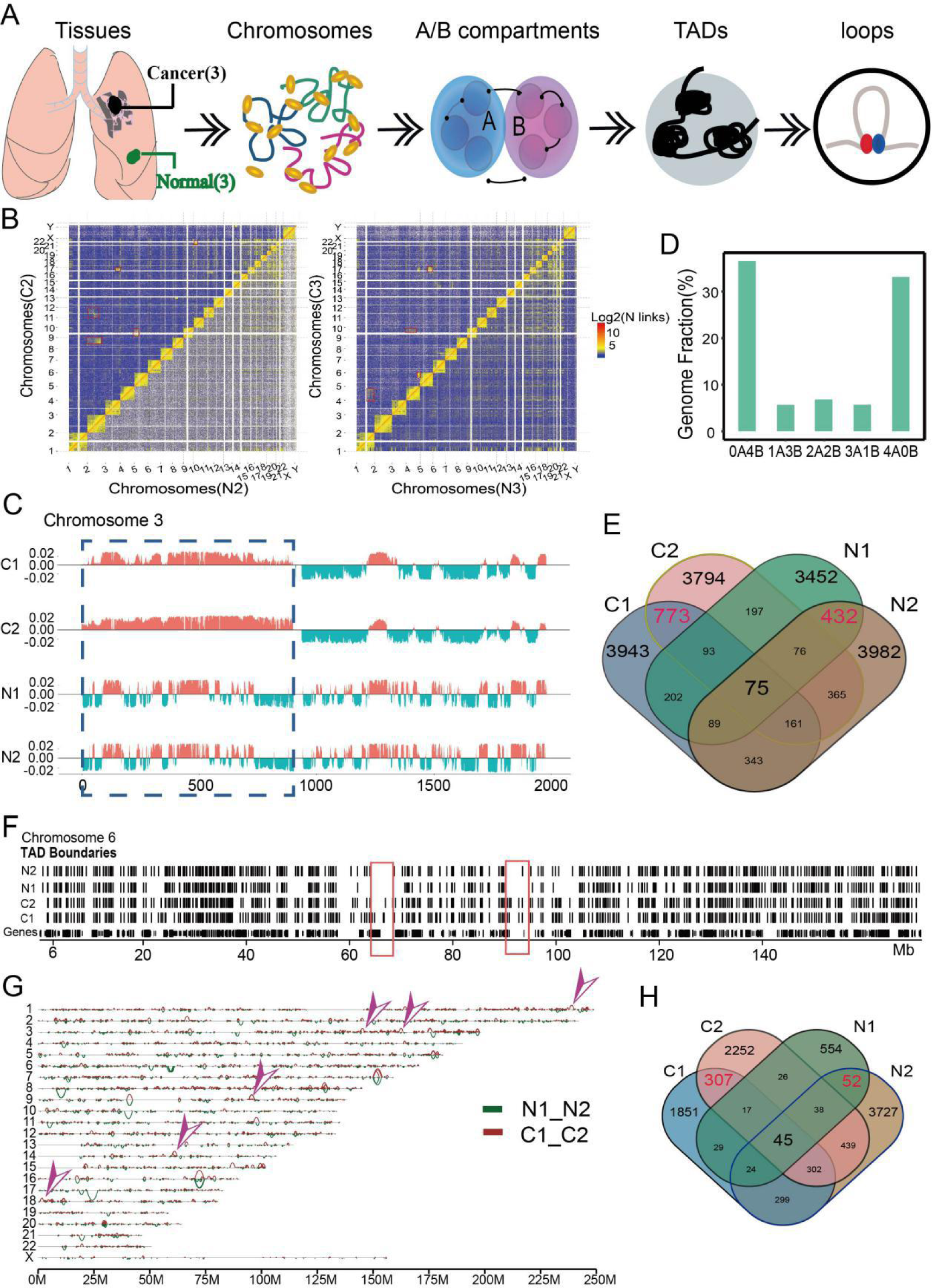
Global Features of 3D Genome Organization in 2 LUAD tumor and normal tissues. (*A*) Sample information and analysis workflow of this study. (*B*) Interaction heatmap of all genome in tumor and normal tissue, and obviously translocation sites were signed by red box. (*C*) Genome browser snapshot showing compartment A/B patterns (PC1 value) across chromosome 3 in 4 samples. Positive PC1 in red corresponds to compartment A, and negative PC1 in green corresponds to compartment B. (*D*) Bar plots showing the degree of conservation of A/B compartment labels of 4 tissues. The y axis is the fraction of the genome conserved by the 5 possible combinations of compartment A/B designations. The label below each bar represents the composition of the compartment designations. For example, ‘‘1A/3B’’ represents the genomic region where 1 samples exhibit a compartment A label and the other three samples exhibit a compartment B label. (*E*) Venn plot showing the overlap of identified TADs in 4 samples. The red numbers represent the overlapped TADs in tumor and normal tissues. (*F*) Genome browser snapshot showing topological domain boundaries across chromosome 6 in 4 samples. Boundaries are identified at 40-kb bin resolution, and cancer specific genome regions were signed by red box. (*G*) Genome browser snapshot showing chromatin loop across all chromosomes in tumor and normal tissues, and tumor specific loop was marked by purple arrow. (*H*) Venn plot showing the overlap of identified loops in 4 samples. The red numbers represent the overlapped loops in tumor and normal tissues.

Chromosomal loci interactions could lead to genome rearrangement which induced spatial structure variation in many tumor-related genes and contributed to occurrence of tumors (Wu P et al. 2017). In this study, the ration of HIC reads number observed in any two chromosomes to expected interaction reads was defined as interaction frequency between chromosomes, the results revealed that interaction frequency between small gene-rich chromosomes (Chr16-22) was lower in tumor tissues compared with normal tissues (Fig. 1B, S1A), while increased interaction was observed in other chromosomes such as chr2 and chr3 in patient 1, chrY and chr14 in patient 2 (Fig. 1B). And interaction frequency decreased rapidly with the increased chromosome distances (Fig. S1B).

Genome interaction contained two major modes, representing open (type A) and closed (type B) genome compartments (Lieberman-Aiden E et al. 2009). Here, we identified multiple A/B compartments based on PCA value of the correlation coefficient matrix from chromatin interaction by HiTC (v1.24.0) software at 100kb resolution (Servant N et al. 2012). The most obvious A/B compartment conversions were found in Chr3 in tumor tissues compared with normal tissues (Fig. 1C), followed by Chr4 and 12 (Fig. S1C).

As previously reported for cultured human cells (Dixon JR et al. 2015), we observed substantial compartment A/B switching across primary tissues (Fig. 1C, D), finding that 23.67 % of the genome is dynamically compartmentalized in different tissues. These data also underscore the significant degree of compartment variation across the genome, revealing that as much as 10.1 % of the compartment related genome is variant (Fig. 1C).

We then analyzed differences of TAD structures between tumor and normal tissues, which is the basic unit of gene expression and is important in forming independent regions of gene regulation (Crane E et al. 2015). TadLib (Franke M et al. 2016) and HOMER (Wang XT et al. 2017) were conducted to identify TAD structures at 20-kb bin resolution (Table S2), which revealed total of 17175 and 15629 TADs in tumor and normal groups, respectively (Fig. S1D), but the domain size had no obvious difference in two groups (Fig. S1E). Total of 5071 unique TADs were identified in LUAD tissues (Fig. 1E), although majority of these TADs were personal specific, we still identified TAD boundaries which were overlapped in two tumor tissues, which might play important roles in tumorogenesis (Fig. 1F, red boxes).

Finally we analyzed the loop structures in LUAD tumor and normal tissues. Chromatin loops occurred in TADs or sub TADs and formed enhancer-promoter interactions, which played important roles in tumoregnesis by increasing expression of oncogenes (Ji X et al. 2016). Here we performed HICCUP software at 10-kb resolution to process HIC data and identify significant interaction sites, the number and mean length of loops were listed in Table S4, showing that more but smaller loop structures were identified in cancer tissues by compared with normal tissues (Fig. 1G). Total of 45 loops were observed in all samples and 307 loops were identified specific in cancer tissues (Fig. 1H). Three types of loops were identified in our study, including enhancer-enhancer (E-E), enhancer-promoter (E-P) and promoter-promoter (P-P), which increased dramatically in tumor tissues (Figure S1E-S1G), suggesting important roles of these loops in tumor progression.

Our results revealed multiple structure variations, including A/B compartment, TADs and loops in three LUAD tumor tissues, which might play important roles in tumor process.

### Conversions of A/B Compartment concentrated mainly on chr3 in tumor tissues

In breast cancer, A/B compartment structures in the chromosomes of tumor contained many genes, which related closely to tumor-related pathways such as WNT and TCF/LEF (Barutcu AR et al. 2015). Here we extensively analyzed A/B compartment conversion in LUAD tumors and performed hierarchical clustering and visualization of these converted AB compartments (Table S3). The results indicated that A/B conversion occurred in every chromosome of all samples (Fig. 2A), while A→B was the main type transition in tumor tissues (Fig. S2A, B), which suggested that multiple genes lost transcriptional activity and prevented the body from performing normal functions and promoted cancer development. In all chromosomes, we noticed that most A/B conversion events were occurred in Chr3 (580/1728, 33.6%) and the dominant type was B→A conversion (355/580) (Fig. 2A).

**Figure 2.**
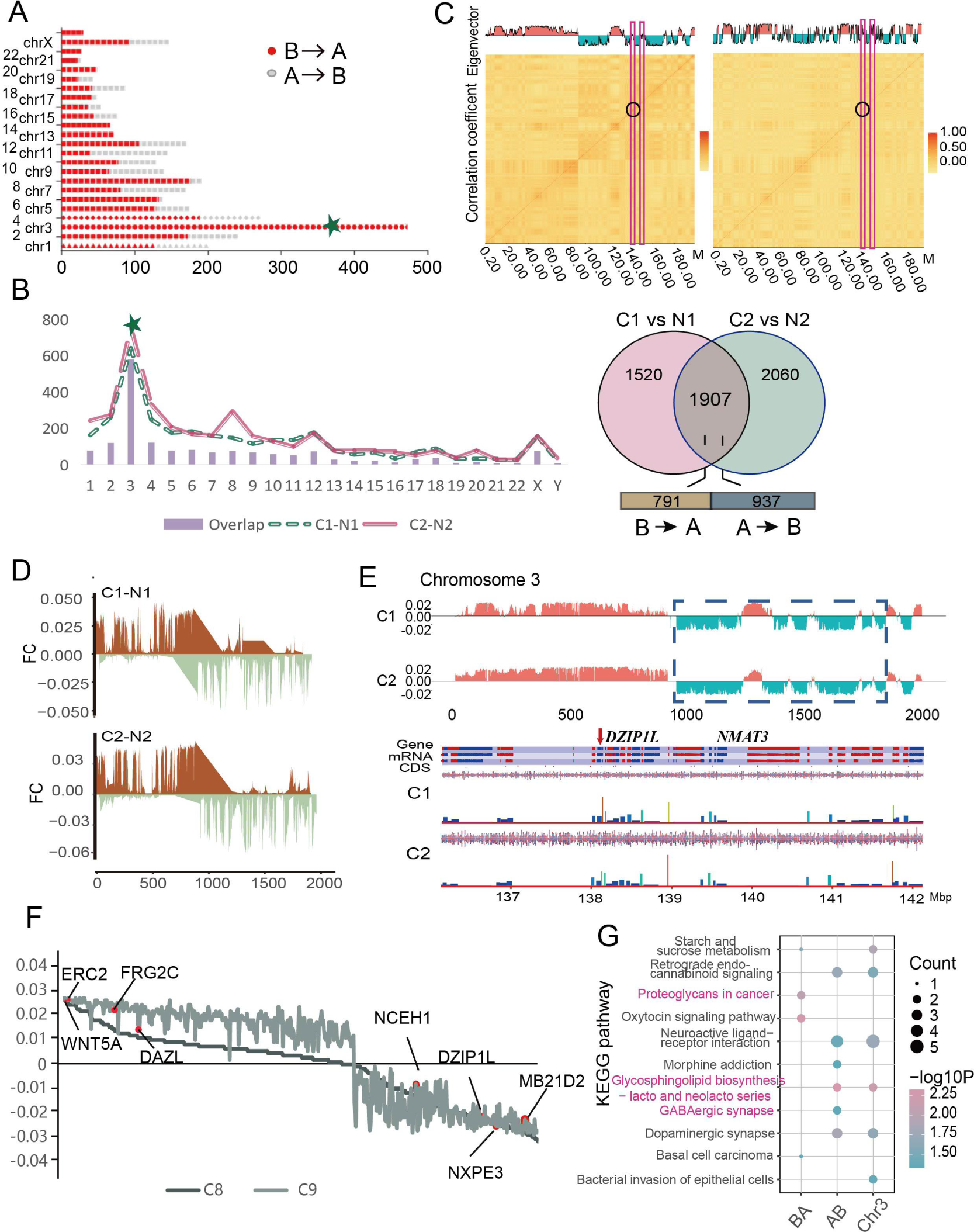
Conversions of A/B Compartment concentrated mainly on chr3 in tumor tissues. (*A*) Distribution map of the number in two compartment conversion types (A→B and B→A) of each chromosome in tumor tissue. The green five-pointed star signing the chr3, which contain the most different compartments. (*B*) Statistics distribution chart of different and overlapped compartment in each chromosome and the of 2 tumor tissues. Venn plot showing the number of overlapped and sample specific compartments and the number of different conversion type in the overlapped compartments. (*C*) Heatmap of A/B compartment interaction in tumor and normal tissue.The horizontal and vertical coordinates of the figure below are the chromosome positions, and the darker the color, the higher the correlation. (*D*) Distribution map of difference between tumor and normal tissue in each patient based on PC1 value across chromosome 3. (*E*) Gene annotation on the regions with compartment conversion of chr3by the SeqMonk Course Manual solfware. (*F*) Distribution map of different compartment related genes in each sample based on the PC1 value, and some important genes were marker on plot. (*G*) KEGG pathway analysis of different conversion of compartment statue on chr3in tumor.

We then analyzed the A/B switch in each tumor tissues and results also showed that Chr3 exhibited the most A/B compartment switches, total of 1907 A/B compartment variations were found in tumor tissues while 580 (30.4%) located on Chr3 (Fig. 2B, S2C). The association analysis between A/B compartment on Chr3 and chromosome genome correlation indicated that correlation intensity of A→B in the cancer tissue also higher than normal tissues (Fig. 2C), which meant the formation of new chromatin structures and promoted tumoregenesis.

Next, we conducted subtraction between compartments of each pair of samples based on PC1 value on whole genome and Chr3 to assess the degree of structural variation of the compartment, the region of Chr3 with the most frequent and highest degree variation was clearly divided into two regions, the first 12.5Mb tends to be converted from A to B, and the rest tend to be converted from B to A (Fig. 2D, S2D), and these regions contained most important genes in *ERC2, WNT5A* and *NMAT3* (Fig. 2E).

Finally we further visualized the 89 compartments identified in tumor tissues, which revealed most important tumor-related genes, including *ERC2, FRG2C*, DAZL, and *MB21D2*, et al (Fig. 2F). Pathway analysis indicated that A→B conversion enriched in glycosphingolipid biosynthesis − lacto and neolacto series and dopaminergic synapse pathways (Fig. 2G). We also annotated all genes of A/B compartment and revealed cancer associated signaling pathways, including PI3K−Akt, MAPK, Ras and Rap1 (Fig. S2E). In general, we demonstrated that A→B was the major conversion type in LUAD tumors and related closely to important signal pathways in tumor process.

### Distinct TAD boundaries variations were identified in LUAD

Occurrence of TAD boundaries was a common characteristic and more than 85% TAD were conserved in cells, however, tumor cells still harbored distinct specific TAD boundaries, involving tumor-related genes such as *GATA1, SOX9, MYC* and might contribute to cancer development (Spielmann M et al. 2018). Therefore we compared the specific TAD structures and boundaries between tumor and normal tissues at 40-kb resolution to observe the specific TAD structure in LUAD and the functional related genes by DI (Directionality index score) delta value (Wang XT et al. 2017). Total of 1500 different TADs were detected in this study, tumor and normal tissues harbored 650 and 850 TADs respectively (Fig. 3A). In TADs distribution analysis in different chromosome, we found that chr2 contained highest number of TADs (149) and chr1 (109), chr3 (107), chr4 (96) and chr8 (104) also had more TADs than other chromosomes (Figure 3A). We then analyzed the TADs in tumor tissues and we identified 446 and 420 TADs in C1 and C2 tumor tissues, 216 TADs were overlapped in both tumor tissues (Fig. 3B). In chromosome distribution analysis we found that majority of overlapped TADs were concentrated in chr1 (19), chr2 (15) and chr3 (17) (Fig. 3C), which revealed the importance of the three chromosomes in TAD formation.

**Figure 3.**
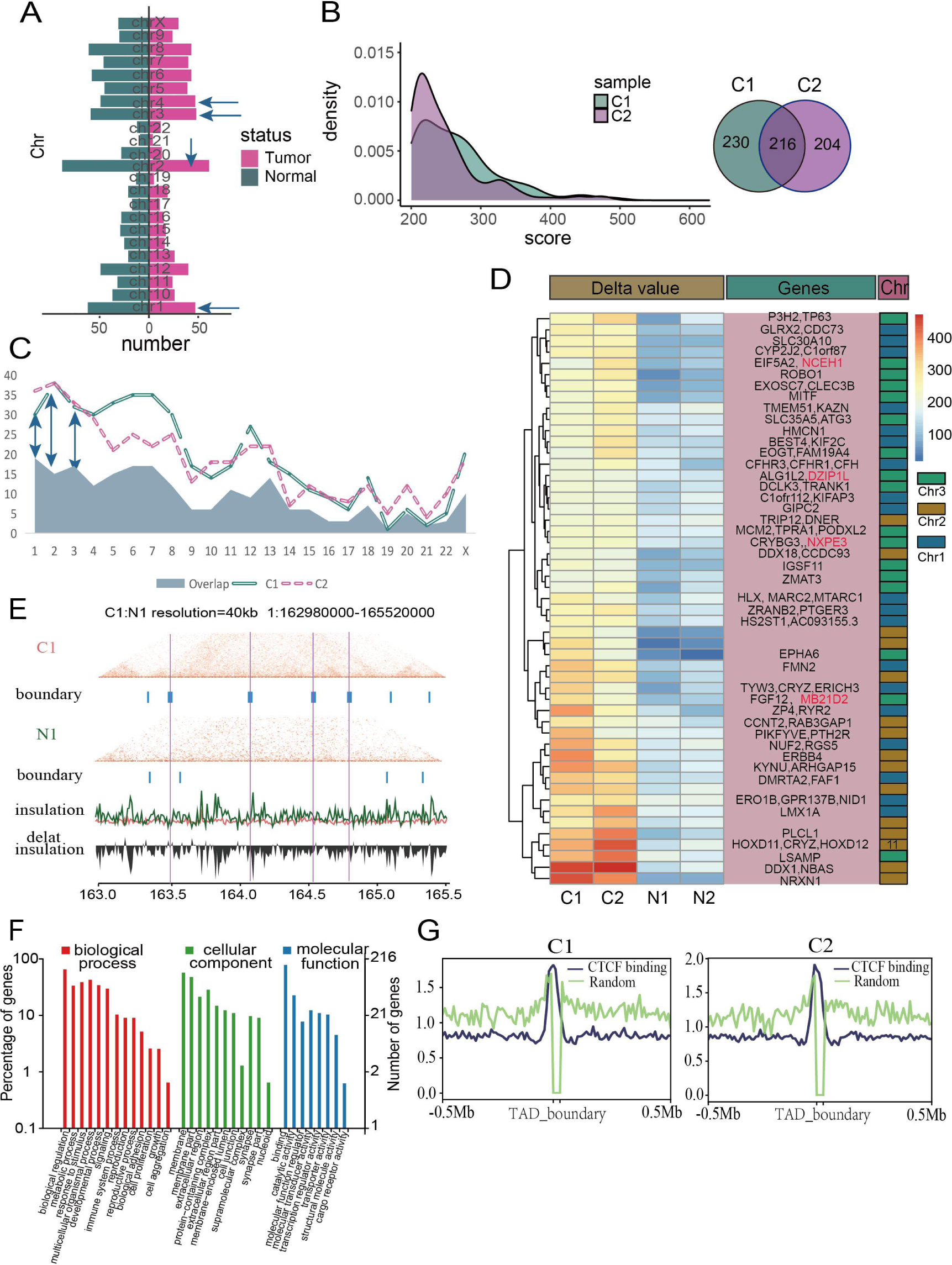
Distinct TAD boundaries variations were identified in LUAD. (*A*) Barplot showing the number of tumor and normal tissue specific TADs in each chromosome, and top 4 chromosomes were marked by green arrow. (*B*) Density distribution of the delta score value in 2 tumor tissues, and venn plot showing the number of sample specific and overlapped in tumor tissues. (*C*) Statistics distribution chart of different and overlapped TADs in each chromosome and the of 2 tumor tissues. (*D*) Heat map of tumor specific TADs of chromosome1,2, and 3based on delta score value, and the related genes was annotated in the pink box on the right, *NCEH1,DZIP1L,NXPE3*, and *MB21D2* were signed by red color. (*E*) Hi-C interaction matrix of a region (chr1: 162980000-165520000) in patient1 shows the difference between tumor and normal tissue with the TAD boundary.Top: Hi-C interaction matrix and TAD boundaries (vertical bars), middle: insulation scores, bottom: delta insulation score. The pink line label the tumor specific TAD boundary. (*F*) GO enrichment analysis of different TADs related genes. (*G*) CTCF binding distribution in tumor based on the public Chip-seq data.

Next, we analyzed the important genes involved in overlapped TADs in chr1, 2 and 3, heatmap results revealed multiple tumor-related genes in TAD boundaries, including *C1orf87, CFHR3* and *RGS5* in chr1, *HOXD11, DDX1* and *NRXN1* in chr2, *TP63, ATG3, LSAMP* and *FGF12* in chr3, the most interesting observation in TADs was that we also found five tumor-related genes, *SYT16, NCEH1, DZIP1L, NXPE3* and *MB21D2*, which were former identified in A→B compartment (Fig. 3D), suggesting the important role of the 5 genes in specific structure variation in lung cancer. In extensive analysis by boundary insulation, we found that significant boundaries were detected in tumor tissues which clearly indicated the formation of novel TADs (Fig. 3E, S2A).

Then we performed ClusterProfiler R package to investigate the important pathways of tumor TADs and we found the variations of pathways concentrated in biological process, cellular component and molecular function, including multiple pathways such as biological adhesion, cell junction, transporter activity, cargo receptor activity and reproductive process which played critical roles in tumor process and metastasis (Fig. 3F). In KEGG pathway analysis, we also identified important tumor-related pathway including cell adhesion molecules, choline metabolism in cancer, as well as hippo, wnt and mTOR signaling pathways (Fig. S3B).

Finally we explored the CTCF(CCCTC-binding factor) in tumor tissues, CTCF is a exceptional transcription factor, and acted as important master in organization of genome into TADs, which was critical for coordinated transcriptional regulation, chromatin states, DNA replication and establishing TAD boundaries (Katoh Y et al. 2006; Wang Y et al. 2020; Barutcu AR et al. 2018), we used the Chip-seq from study of ENCODE Project Consortium (2012) to analyze the CTCF distribution of TAD boundary in important genes, which showed that CTCF had a higher reads ratio value in TAD boundaries (Fig. 3G, S3C).

Taken together, we found 216 tumor-specific TADs in LUAD samples and further identified five important genes (*SYT16, NCEH1, DZIP1L, NXPE3* and *MB21D2*), which were consistent with A/B compartment results.

### Specific chromatin enhancer-promoter loops were identified in LUAD tumor tissues

The chromatin loops increased formation of active chromatin hubs composed by interactions of multiple enhancers and promoters (Ji X et al. 2016), and previous studies indicated that chromatin loops could override endogenous gene expression programme and provide a possible therapeutic approach in disease treatment such as sickle cell anaemia (Deng W et al. 2014; Deng W et al. 2012; Zheng H and Xie W 2019). Some strong interaction signal points at off-diagonal positions was found in the whole genome heat map in Figure 1B, which suggested the existence of loop structures. Here, we explored the loop structures at the 10kb resolution and FDR <= 0.01 by the HICCUP software.

To compare the loop structures between tumor and normal tissues, review’ method was performed (Zhang Y et al. 2019) to analyze all samples. Total of 2344 different chromatin loops were identified, 1357 loops existed specific in tumor tissues (Fig. 4A, Table S4) and the mean length was lower than normal tissues (Fig. 4A). All loops were counted and visualized in all chromosomes and the results revealed that most chromatin loops (312) concentrated in chr3, followed by chr1 (230), chr2 (155) and chr8 (220) (Fig. 4B, S4A). And we revealed that C1 and C2 tumor tissues harbored 956 and 1012 loops respectively, total of 131 loops were overlapped in both tumors (Fig. 4C, inner Venn), majority of overlapped loops dominated in chr3 (Fig. 4C), which indicated the important role of chr3 in formation of chromatin loops.

**Figure 4.**
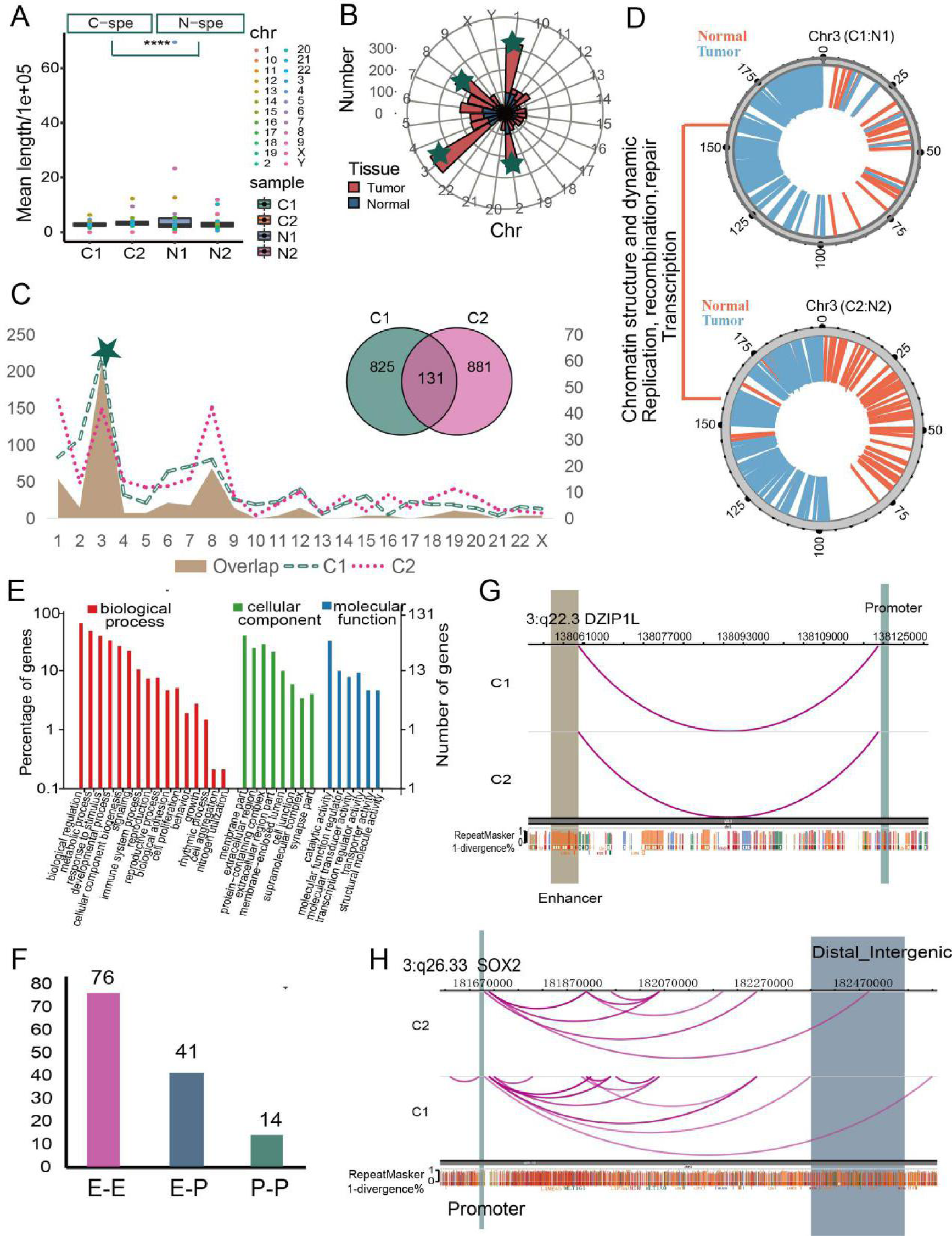
Specific chromatin enhancer-promoter loops were identified in LUAD tumor tissues. (*A*) Boxplot showing the mean length of tumor and normal specific loops in4 samples. (*B*) Polar diagram showing the number of tumor and normal tissue specific loops in each chromosome, and top4 chromosome were marked by green five-pointed star. (*C*) Statistics distribution chart of different and overlapped loops in each chromosome and the of 2 tumor tissues, and green five-pointed star signed the chr3, which has most number of different loops. Venn plot showing the number of overlapped and sample specific loops. (*D*) Distribution of tumor and normal specific loops on genome of chromosome 3, and showing the function of these genome regions. (*E*) GO enrichment analysis of the tumor specific loops related genes. (*F*) Barplot showing the number of loop types of 131 overlapped loops in 2 tumor tissues. (*G*) Genome browser snapshot of chromatin loop with the annotation of the end anchors at *DZIP1L* locus, the brown bar represent the enhancer domain of gene, green bar represent the promoter domain and the blue bar is the distal enhancers. (*H*) The same visualization with (G) at the *SOX2* locus.

Next, HiCPlotter was conducted to compare the differences of important genome regions in formation of chromatin loops between tumor and normal tissues and we found that tumor-specific loops were composed by *SIDT1, HMCES, NMNAT3*, and *GPR87*, which all related closely to tumor process (Fig. 4D). We then visualized all cancer specific loop structures in chr3 and annotated these chromosome distinct genes with GO enrichment, which revealed that pathways of these loops concentrated in biological process, such as immune system process, cell proliferation and biological adhesion, cellular components including cell junction, protein-containing complex and supramolecular complex, molecular function including molecular transducer activity, transcription regulator activity and transporter activity (Fig. 4E), majority of pathways (Fig. S4C) were tumor-related and confirmed the important of loops in tumoregnesis and chr3 was the main source of loops structures in tumor.

Enhancer-enhancer (E-E), enhancer-promoter (E-P) and promoter-promoter (P-P) were the three major types of chromatin loops and promoter-enhancer loops played important regulatory roles in occurrence and development of diseases (Deng W et al. 2012). In this study we obtained 76 E-E, 41 E-P and 14 P-P loops (Fig. 4E), all enrolled genes were listed in Table S5. The five genes (*SYT16, NCEH1, DZIP1L, NXPE3* and *MB21D2*) formerly identified in A/B compartment and TADs, were also found in E-P loops in chr3. Among five genes, the anchor site at two ends of the loop, which located on q22.3 of chr3, was formed by *DZIP1L* promoter and promoter, another chromatin loop was formed by a *MB21D2* enhancer, we also observed physical interaction between more than three distal enhancers and the promoter region of the gene *SOX2*, which on chromosome 3, a known oncogene that stimulates cellular migration and anchorage-independent growth (Fig. 4F, S4D). The results confirmed the importance of chr3 in formation of chromatin loops.

### *SYT16, NCEH1, NXPE3, MB21D2, DZIP1L* were the most important genes in advanced structure variations in LUAD

3D organization of cancer genome induces chromosome instability and leads to occurrence of tumors (Barutcu AR et al. 2015).The spatial organization of genome also shapes alteration of cancer genome (Fudenberg G et al. 2011). Our former analysis confirmed that cancer-specific advanced structure variations, including A/B compartment, TADs and chromatin loops, were observed in LUAD tumor tissues. To further identify important genes which may involve in formation of advanced structure variation, we put A/B compartment, TADs and chromatin loops altogether to find the overlap molecules in all three structure variations and the results revealed five genes *SYT16, NCEH1, NXPE3, MB21D2*, and *DZIP1L* (Fig. 5A). In all five genes, *NCEH1, DZIP1L, NXPE3* and *MB21D2*, located on p arm of chr3 and *SYT16* located on chr14 (Fig. 5B, C). *SYT16* was related to calcium ion binding and protein heterodimerization activity, which also acted as prognostic biomarker in LGG (Lower-grade glioma) (Chen J et al. 2020). *NCEH1* expressed in human atheromatous lesions and played a critical role in the hydrolysis of CE in human macrophage foam cells, thereby contributing to the initial part of reverse cholesterol transport in human atherosclerosis (Igarashi M et al. 2010). *NXPE3* was significantly hypermethylated and down regulated in HCC (hepatocellular carcinoma) tumors, and epigenetic silencing of the gene may be associated with occurrence of HCC (Yamada N et al. 2016). Mutated *MB21D2* was found in LUAD tumor tissues and suggested an oncogenic role and could be a potent neoantigen liked other passenger mutation (Buisson R et al. 2019). *DZIP1L* functioned as important molecule in Hedgehog signaling pathway and often activated in gastric cancer (Katoh Y and Katoh M 2006).

**Figure 5.**
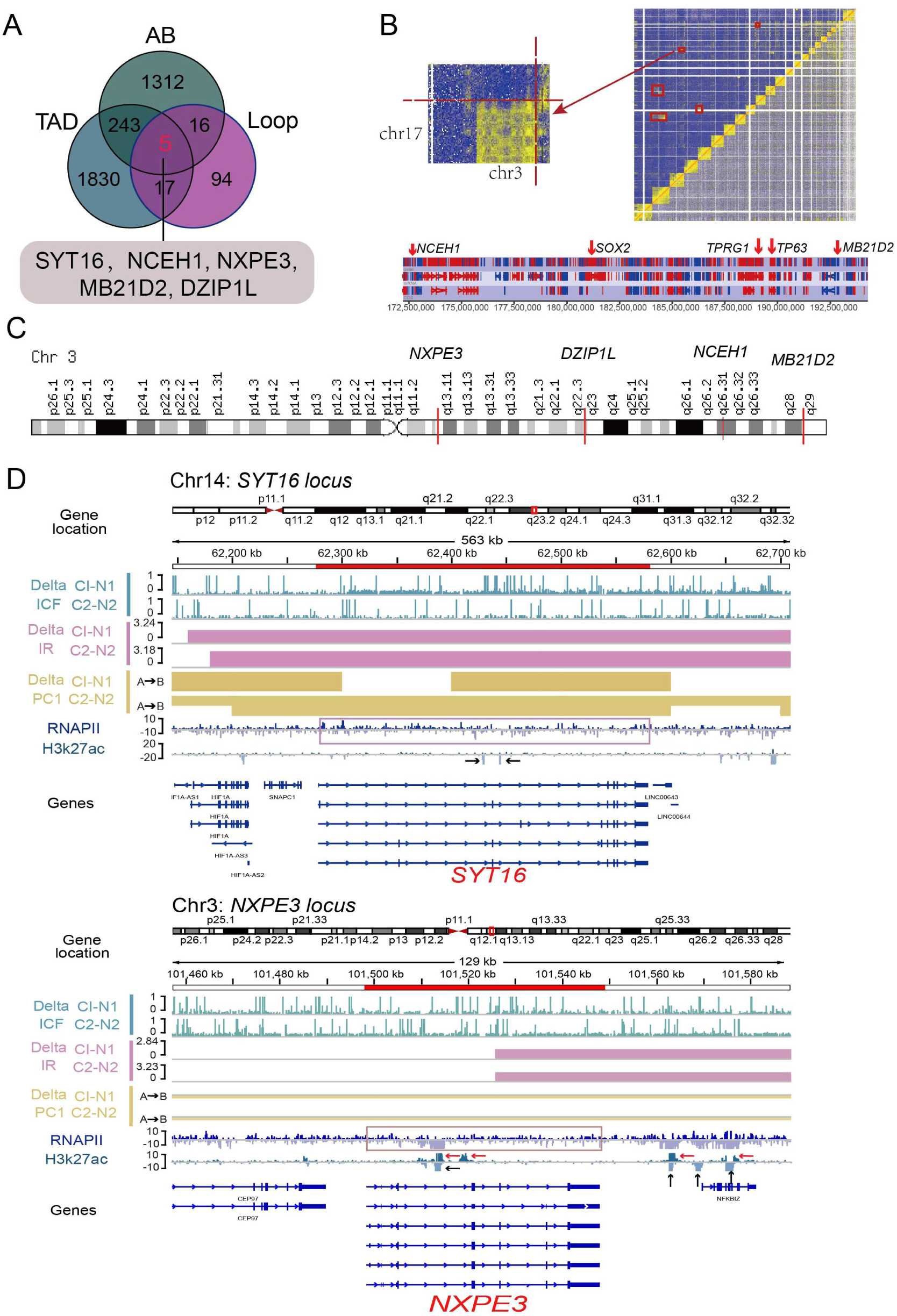
5 most important genes in advanced structure variations in LUAD. (*A*) Venn plot showing the related gene number distribution of tumor specific chromatin structures (Compartment, TAD, loop), and 5 genes(red number)were found in each resolution chromatin structure. (*B*) Heatmap and partial heatmap of tumor showing a balanced translocation between chromosomes 3 and 2. Heatmaps were coloured by the number of interactions with the colour gradient scaled linearly from 5 (blue) to 10 (red). The bottom plot showing read counts for amplified regions on chromosomes 3. The high peaks show a significantly higher number of reads than in the surrounding regions.*NCEH1, SOX2, TPRG1, TP63* and *MB21D2* genes are labelled. (*C*) Distribution of the position of 5geneson chromosome3 genome, and 4 genes (*NXPE3, DZIP1L, NCEH1, MB21D2*) are labelled. (*D*) Genome browser tracks of ChIP-seq (RNAPII/H3K27ac), compartments changes (Delta PC1), TAD changes (Delta IR) and loop changes (Delta ICF) in tumor compared with tumor tissue at the *SYT16* (top) and *NXPE3* (bottom) genes locus.

To explore the association between advanced chromatin structure (compartment,TAD, and loop) variations and transcription and regulation of the 5 genes, we used the public Chip-seq data (RNAPII and H3k27ac) from previous studies (Sammons MA et al. 2015; Kurppa KJ et al. 2020) to evaluation (Fig. 5D, S5), the RNAPII can converts the DNA sequence of a single gene into the multiple alternative functional transcript isoforms and H3k27ac is an active enhancer marker. At the *SYT16* locus on chromosome 14, the regions of gene showed increased decompaction (stronger interaction of chromosomes and higher insulation score) and A-to-B compartment switching with the presence of RNAPII in tumor tissue, but the deletion of H3k27ac signal. The same phenomenon was observed at the *DZIP1L* and *NCEH1* locus on chromosome 3 (Fig. S5A, B). At the *NXPE3* locus, expect gene region, the region downstream of gene also showed the same chromatin structure variations in tumor, and the lower expression of RNAPII with the deletion of existing H3k27ac signals and generation of novel H3k27ac signals. The similar phenomenon was observed at the *MB21D2* locus (Fig. S5C). Results suggesting that stronger activity of RNAPII while A-to-B compartment switching and may generate new regulation site.

Taken together, 5 genes was found have variations in the AB, TAD and loop structure of genome with the change of RNAPII and H3k27ac signals in tumor, implicates a potential role for these genes in facilitating the occurrence and development of LUAD.

## Discussion

Chromatin instability played important roles in occurrence of malignant tumors, advanced chromatin structure of genome, such as A/B compartment, topological associated domains (TADs) and chromatin loops has now become one of the hottest topics in tumor fields (Hui Z et al. 2019; Vera BK et al. 2017; Anne-Laure V et al. 2016). Earlier studies about 3D genome with Hi-C technology in cancer largely limited to cultured cells or a small collection of primary cancer cell types, such as myeloma and prostate cancer (Pengze W et al. 2017; Phillippa CT et al. 2016; Suhn KR et al. 2019), which ignored the impact of tumor microenvironment and contributed poor manifestation to the true pathological characteristics of malignant tumors, and previous study showed that there was a genomic inconsistency between subcultured cells (U87MG glioma cell line) and primary tumor cells (glioblastoma) (Marie A et al. 2016).

In our study, it was the first time to apply Hi-C technology to detect clinical lung adenocarcinoma samples and conducted in-depth analysis about the variations of advanced chromatin structure in LUAD compared with normal tissues. We observed multiple advanced structural variations (A/B compartment, TADs and loops) in tumor tissues. Our results confirmed the dominant role of chromosome 3 in occurrence of lung tumor structural variations, which harbored 820/5308 (15.4%) A/B compartment, 107/650 (16.5%) TADs and 312/1357 (23%) chromatin loops. Study in lung cancer revealed that gain/amplification at q and loss/deletion at p arms of chromosome 3 were the most frequent and early events in lung cancer, encoded many candidate driver genes, contain *PIK3CA* and *TP73L*, participate in cancer associated functions, such as cell proliferation, cell adhesion, apoptosis, RAS signaling, and oncogenic transformation(Qian J et al. 2008). Total or part copy deletions in p arm of chromosome 3 related closely to development of clear cell renal carcinoma, and code many olfactory receptors as well as chemokine receptors that aid inflammatory processes, such as *CCR5* and *VHL* (Brunelli M et al. 2011). Our study also identified many tumor-associated genes, such as *ERC2, WNT5A* in compartment, *TP63, MITF* in TAD structure and *SOX2, NMNAT3* at loop scale and these genes are mainly participated in cell proliferation, transporter and regulation functions and MAPK, PI3K-AKT, Ras, Wnt, Ras1 pathways.

A→B compartment conversions were frequently observed in malignant tumors cells (Barutcu AR et al. 2015), but it was still remained unknown in lung cancer. In our study, 26.7% of genome compartment variation and a higher close compartmentalization during the LUAD development related closely to accessibility of transcription factors or other regulatory proteins, such as *ZDBF2, EDIL3* and *HAPLN1*, which were particularly important for certain subsets of genes (Dixon JR et al. 2015). Our result showed that the genes with variation in compartment status were most notably related to extracellular region and participated in cell proliferation, biological adhesion in cancer, as well as molecular carrier activity, hijacked molecular function and transcription regulator activity.

Besides A→B compartment, TADs and chromatin loops also played important roles in tumor development (Malte S et al. 2018; Wulan D et al. 2014; Wulan D et al. 2012; Hui Z et al. 2019). Our study also unveiled multiple TADs and loops in LUAD tumor tissues, the interesting observation was that not all changed topologically domains (TADs) were functional consequences of genes (Figure 3D), non-coding regions in of TADs were also identified in lung tumors, suggesting the regulatory roles of these regions but still need further investigation. We also observed some novel TADs (Figure 3E and S3A-S3B) in tumor tissue by compared with normal sample, the formation of these structures may due to the duplications spanning a TAD boundary (Martin F et al. 2016).

Furthermore, in our study many E-P type loops were identified in cancer tissues, containing tumor associated genes at loop anchor points, such as *SOX2,TPRG1*, which were reported exist somatic variation and were key upregulated factors in lung cancer, involved in tumor progression. Previous study identified enrichment of SNPs for different sets of diseases in each set of group-specific loops by the analysis similar to GO enrichment (Grubert F et al. 2020), so intersecting the variants in promoter or enhancer regions of these oncogenes with tumor-specific chromatin loops in our study may help to explain how such sequence variation leads to LUAD. We also observed obvious individuality (Figure 4C and S4B) at the loop scale, but the mechanism for this phenomenon remained further exploration.

The most important observation in our study was that 5 genes (*SYT16, NCEH1, NXPE3, MB21D2* and *DZIP1L*) were identified in the variations of both A/B compartment, TADs and chromatin loops in tumor tissues. Among 5 genes, 4 (*NCEH1, NXPE3, MB21D2*and*DZIP1L*) of the 5 genes on the q arm of chromosome 3, which further confirmed the important role of chromosome 3 in LUAD. *NCEH1* is a tumor-associated enzyme, involved in the general processes of tumor progression and may serve as a marker for malignant potential (Igarashi M et al. 2010), *NXPE3*, the epigenetic silencing of the gene may be associated with occurrence of HCC (hepatocellular carcinoma) (Nobuhisa Y et al. 2016), *MB21D2* is correlated with poor survivals in patients with head and neck and endometrial cancer and as a oncogene in LUAD (Buisson R et al. 2019), *DZIP1L* has reported as a second gene involved in ARPKD (Autosomal recessive polycystic kidney disease) pathogenesis (Katoh Y et al. 2016). Another gene, *SYT16*, on the chromosome 14 is also as a prognostic biomarker and correlated with immune infiltrates in glioma (Chen J et al. 2020). Take together, the 5 genes found in our study may play a key role in the development of LUAD and also revealed the importance of chromosome 3 in chromatin structure.

Genomic instability in cancer is not a single phenomenon. Instead, many different mutational processes can act to restructure the genome and, in doing so, generate a notably flexible array of possible structures. In this study, we did a comprehensive analysis of chromatin structure in LUAD using HIC technology firstly, and found the importance of chromosome 3 and 5 meaningful genes, which can provide reference basis for future research. Although we have found some important chromatin structural variations and genes in lung cancer tissues, we need to further combine other omics data for joint analysis to interpret the genomic structural variations of tumor tissues from more perspectives.

## Methods

### Sample Collection

Two pairs of fresh tumor tissues and adjacent normal tissues were collected from patients with primary lung cancer undergoing surgical resection without neoadjuvant therapy before surgery and smoking history at West China Hospital (WCH), then store samples at −80°C until use and and all patients provided informed written consent.

### Hi-C experiments

Before starting the experiment, make sure the PBS and formaldehyde added to the sample are freshly mixed and incubate 20% SDS at 37°C for 15 min before use. Disrupt the frozen Tissue(20-40 mg) either by coarsely chopping it with a razor blade on a petri dish placed on dry ice or by grinding it with a mortar and pestle in liquid nitrogen. Then Transfer the disrupted tissue samples to a 1.5 mL microcentrifuge tube which contain 1 ml 1X PBS and 40.5µl 37% formaldehyde, then vortexing the tube for 30s to mix and rotating tube at room temperature for 20 min. Pipeting off and discard the supernatant and washing the pellet with 300 µl wash buffer, after the second wash, added to pellet (99 µl Wash Buffer;1 µl 40 mM Calcium Chloride; 25 µl 1 mg/mL Collagenase), then incubate it for 1 hour at 37 ° C in an agitating thermal mixer (1250 rpm). After incubation, add 6.3 µl 20% SDS to the sample, then pipet up and down to break up any large clumps, after briefly vortex to resuspend, transfer the SUPERNATANT to a new 1.5 mL microcentrifuge tube. The Hi-C experiment was performed following the in situ Hi-C protocol (Rao S et al. 2014). Briefly, the cross-linked cells were lysed and digested with HindIII orMboI, filled with biotin-14-dATP, proximately ligated with T4 DNA ligase andreverse crosslinked with 5M sodium chloride. Then the genome DNA was purified,sheared and size-selected. Biotin pull-down was performed to enrich target DNAfragments, followed by standard Illumina library construction, lastly using a TapeStation or Bioanalyzer to verify the size distribution of your size-selected library.

### Quantification And Statistical Analysis

#### Library quality control and sequencing

After the library construction was completed, the library concentration and insert size were detected using Qubit2.0 and Agilent 2100, respectively, and the effective concentration of the library was accurately quantified using Q-PCR method to ensure library quality. Libraries were sequenced on the Illumina NovaSeq 6000 platform at Novogene, Beijing, China. on average each sample generated 313.12 Gb of raw data.

#### Sequencing data quality control

Quality control (filter joints and low-quality reads) for the raw data, and count the total number of clean reads, the total number of clean data bases, GC content, Q20 and Q30 for each sample and using Phred quality evaluation formula(Qphred = −10log10(e)) to reflect base quality value.

#### HiC library quality assessment

We used Hicup_truncater tool of the HICUP software (Babraham Institute) to estimate the proportion of Truncated reads in the all reads, the ratio should more than 10%. Then the BWA software (Heng Li et al. 2009) (version: 0.7.10-r789, align type: aln) was used to align the double-end data of clean reads to the sequence of the reference genome (ftp://ftp.ensembl.org/pub/release-95/fasta/homo_sapiens/dna), and evaluate the efficiency of the alignment. Then, we used the HIC-Pro v2.10.0 (Nicolas Servant et al. 2015) to analyze the alignment results, identify the valid Interaction Pairs and Invalid Interaction Pairs in the HIC sequencing results, to evaluate the quality of the HIC library. The standard of library quality is: the proportion of Invalid Interaction Pairs cannot exceed 80% (Belton, J.M. et al. 2012). And we also counted the number of interactive fragments Cis/Trans after double-end reads aligned. Finally, the Pearson correlation coefficient was calculated between each biological replicate, to observe the correlation of the biological replicate library.

#### Chromosome location analysis

We calculated the ratio of the number of interaction HIC reads observed between any two chromosomes to the number of interaction reads expected to occur, which is used as the frequency of interactions between chromosomes to predict the strength of interactions between chromosomes in the genome (Yu Zhang, et al. 2012).

#### Resolution analysis

We used the method reported in (Rhie SK et al. 2019) to calculate the resolution of these Hi-C data. Briefly, arrange the number of interactive reads between each bin from high to low according to different size bins, when 80% of the bins are arranged, the bin still covers more than 1000 reads. The size of the smallest bin that meets this condition is the resolution of the HIC library.

#### Interaction analysis

IDE (Interaction decay exponents) is used to describe the relationship between chromosome interaction frequency and interaction distance. In this paper, HIC-Pro v2.10.0 software is used to divide the HIC data according to bin=100kb to calculate the interaction intensity and normalize the data. Then use HICdat (Schmid, MW et al. 2015) to calculate the relationship between the interaction frequency and the linear distance of the genome. The slope of the resulting model is the corresponding IDE value.

#### A/B compartment analysis

##### A/B compartment identification

We used ICE-normalized interaction matrices at 100-kb resolution to detect chromatin compartment types by R-package HiTC (v 1.24.0) (Servant N et al. 2012). Positive or negative values of the first principal component separate chromatin regions into two spatially segregated compartments. The compartment with higher gene density was assigned as A compartments, and the other compartment was assigned as B compartment (Servant N et al. 2012). Then we counted the number of genes on the A/B compartment in each sample.

##### Difference analysis of compartment

The different conversion of compartment in cancer tissue compared with normal tissue is identified as the different AB compartment. We also calculated the FC (Fold Change) value of AB compartment based on PC1 between cancer and normal tissue, to evaluate the degree of conversion of the compartments.

#### TAD analysis

##### TAD identification

We used ICE-normalized interaction matrices at 40-kb resolutionto call TAD by TadLib software (Dixon JR et al. 2015) at bin=20kb. At the same time, IR (Inclustion Ratio) was calculated by HOMER software (Wang XT et al. 2017), indicating the ratio of the interaction between the internal TAD and the interaction of the same length area upstream and downstream from the external TAD, to filter the TAD. The TADs with IR> 1 and the length is more than 5 bins were retained.

##### Difference analysis of TAD boundary

We firstly calculated the DI (Directionality index score) value of the whole genome range of the sample, which is the directional score, then merge the TAD boundaries of all samples, and finally further calculated the DI delta value of each TAD (that is, the average difference of DI in each 4 bins of TAD boundary upstream and downstream). The limma package is used to calculate the difference between cancer and normal tissue. Different TAD boundary was defined as: the difference is significant FDR value <0.01, and the DI delta score value of the two groups of samples is not greater than 200.

#### Loop analysi

##### Loop identification

We used improved HICCUP method (Rao, S et al. 2014, Muzny DM et al. 2006) to identify the loop structures at 10-kb resolution, and the size of bins is 10kb, FDR <= 0.01.

##### Promoter-enhancer prediction

In order to avoid possible deviations in the anchor sites at both ends of the loop, we extend the length of the bin to the left and right of the anchor site by a bin, and intersect the gene transcription start site (upstream 2kb as the promoter region) to obtain each loop annotation information of the anchor site. The anchor site on one side of the loop is located in the promoter region, and the anchor site on the other side is located in the non-promoter region (potential enhancer region, enhancer-like) is called a promoter-enhancer related loop.

##### Difference analysis of loop

Using the method of Yanxiao Zhang, et al (Barutcu AR et al. 2015), firstly, merge the loop results of each sample and de-redundant, then analyze the different loops between tumor and normal tissues based on the original interaction value of each sample, which is calculated for the non-redundant loop, by the edgeR v3.8.6. The different loops are defined as: FDR<1, pvalue<0.05, and FC>1.5. For the different loops, calculate the normalized interaction frequency between the two anchors of each loop, and use R language Visualize.

#### GO and KEGG analysis

GO biological process and pathway enrichment analyses were performed using enrichGO and enrichKEGG from clusterProfiler (Yu, Wang et al. 2012). And the results were visualized with the ggplot2 R package (Wickham 2016).

#### Gene annotation

Firstly, we merge the intervals corresponding to the two samples (for example: merge the IR intervals of C1 and N1), if the sample has no value in this interval, fill it with 0. Then the difference of PCA, IR, and ICF is represented by the corresponding difference between tumor and normal tissue, the difference of RNAPII and H3K27ac between lung cancer and normal cell from public Chip-seq data. Lastly, visualized these genes using IGV solfware.

## Supplementary Material

Table S1: Sequencing result of 2 lung cancer patients by Hi-C

Table S2: Identification number and domain size of TADs in 4 samples

Table S3: Different number of compartment in 2 tumor sample

Table S4: Different number of loop in 2 tumor sample

Table S5: Tumor specific 131 loops related genes list

## Acknowledgments

This work was supported by the National Natural Science Foundation of China (81974363, 81772478).

---

**Figure S1.**
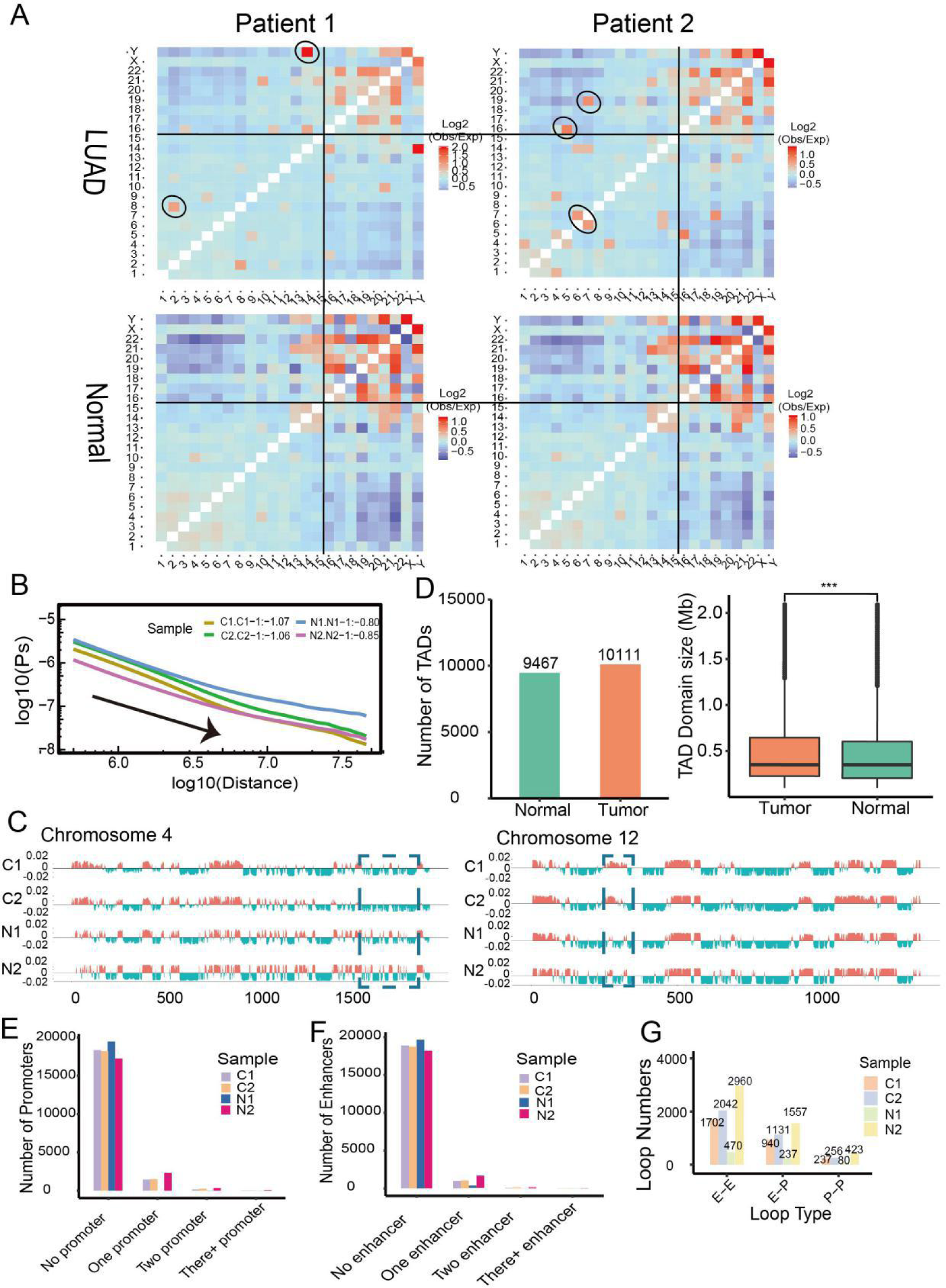
Global Features of 3D Genome Organization in 2 LUAD tumor and normal tissues. (*A*) Heatmap of interaction frequency between chromosomes in patient 1 and patient 2, the small chromosomes (16-22) were separated by black line and tumor specific translocation is signed by black oval. (*B*) Relationship between chromatin interaction frequency and distance in4 samples. (*C*) Genome browser snapshot showing compartment A/B patterns (PC1 value) across chromosome 4 and 12 in 4 samples. (*D*) Barplot and boxplot showing the number of TADs and these TAD domain size intumor and normal tissues, respectively. (*E, F*) Classification chart of anchor sites of loop in 4 samples. (*G*) Classification chart of loop.Based on the anchor site types at both ends of the loop.

**Figure S2.**
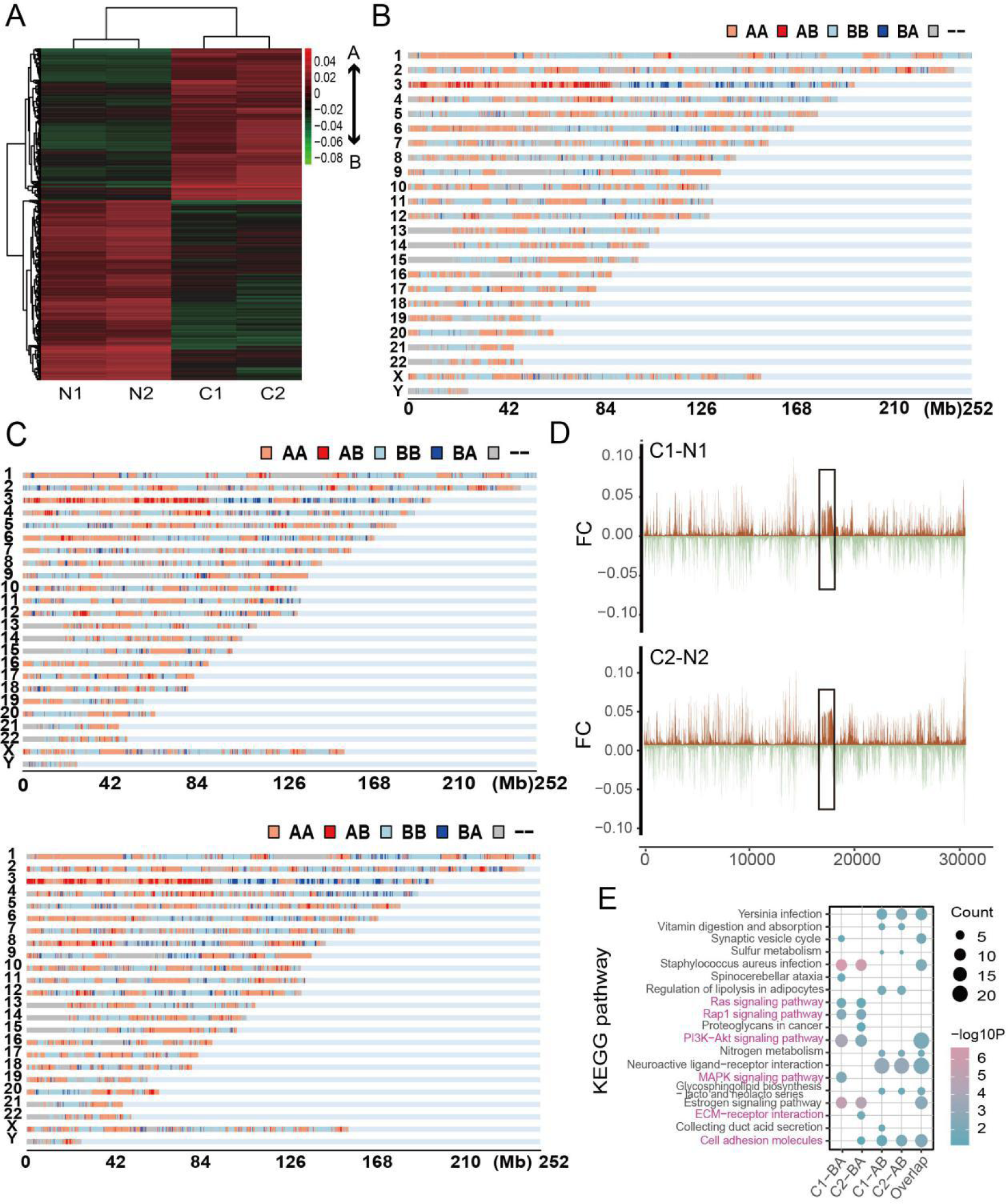
Conversions of A/B Compartment concentrated mainly on chr3 in tumor tissues. (*A*) Heat map of different compartments distribution between tumor and normal tissues based on PC1value. (*B*) Genome browser snapshot showing the different A/B compartment conversion types across all chromosomes between tumor and normal tissues. (*C*) Genome browser snapshot showing the different A/B compartment conversion types across all chromosomes in patient1 and patient2 samples, respectively. (*D*) Distribution map of difference betweenC1 and N1samplebased on PC1 value on the genome,and the region of chr3 is signed by black box. The same distribution map betweenC2 and N2 sample. (*E*) KEGG pathway analysis of different conversion of compartment statue in tumor, and the cancer associated pathway is signed by purple.

**Figure S3.**
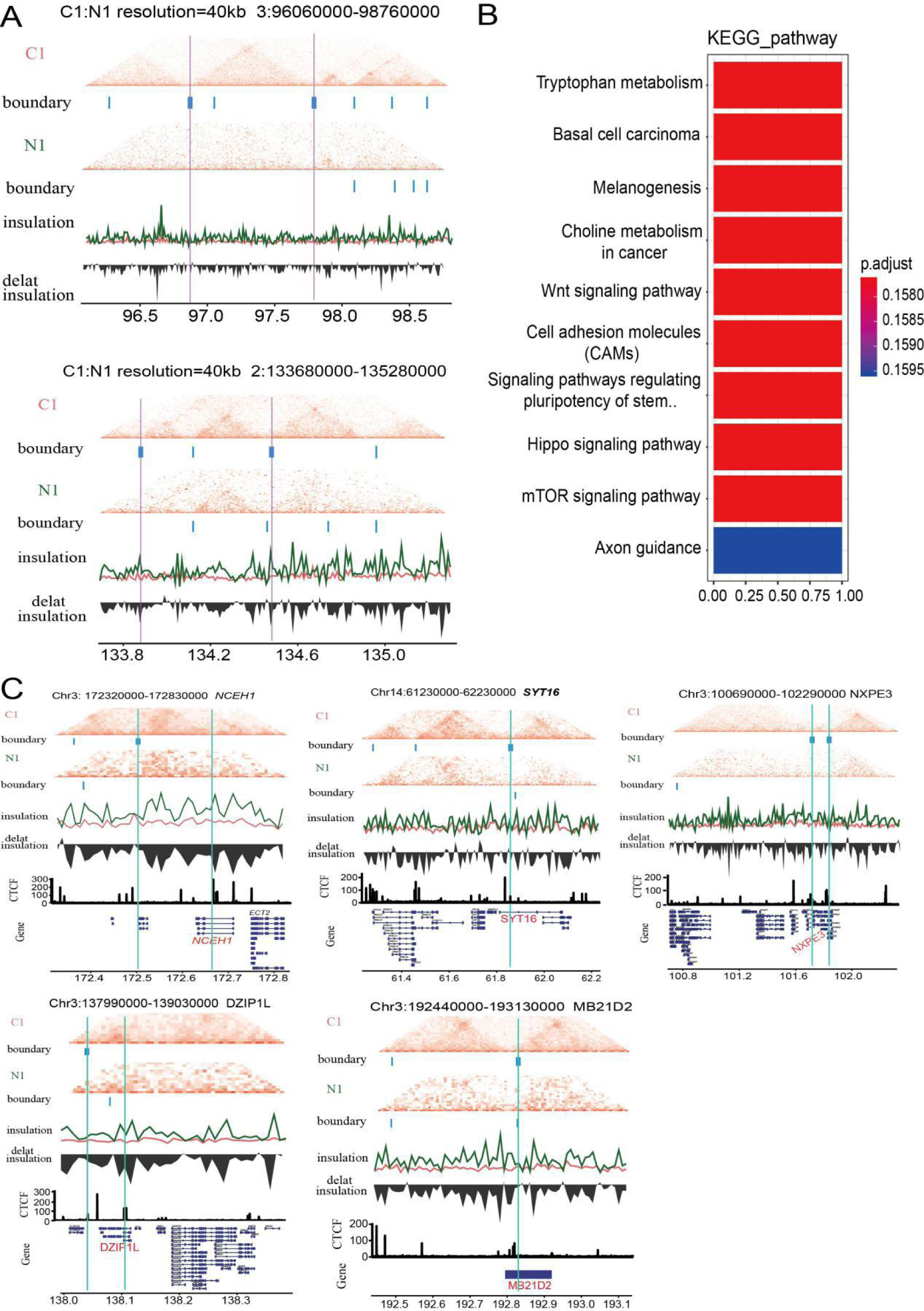
Distinct TAD boundaries variations were identified in LUAD. (*A*) Hi-C interaction matrix of a region (chr1: 133680000-135280000andchr3:96060000-98760000) in patient1 shows the difference between tumor and normal tissue with the TAD boundary. Top: Hi-C interaction matrix and TAD boundaries (vertical bars), middle: insulation scores, bottom:delta insulation score. The pink line label the tumor specific TAD boundary. (*B*) KEGG pathway analysis of different TADs related genes. (*C*) Hi-C interaction matrix of regions contain *NCEH1, DZIP1L, NXPE3, MB21D2* and *SYT16* in patient1 shows the difference between tumor and normal tissue with the TAD boundary and CTCF binding. Top: Hi-C interaction matrix and TAD boundaries (vertical bars), middle: insulation scores and delta insulation score, bottom: CTCF and genes. The green line label the tumor specific TAD boundary and the genes.

**Figure S4.**
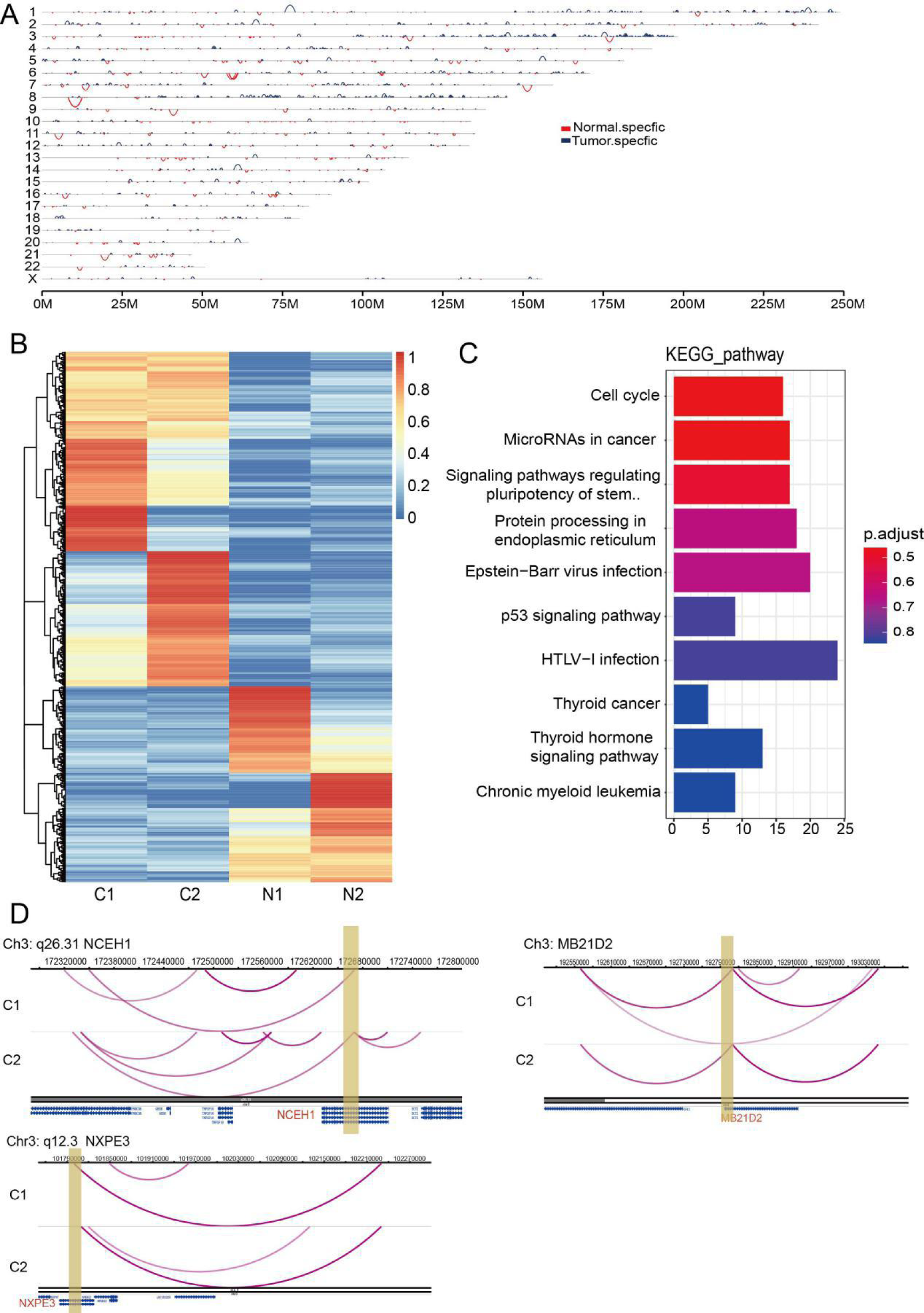
Specific chromatin enhancer-promoter loops were identified in LUAD tumor tissues. (*A*) Genome browser snapshot showing the tumor and normal specific loops, red line show the normal specific and blue show the tumor specific loops. (*B*) Hierarchical clustering heat map showing the difference of tumor and normal samples. (*C*) KEGG pathway analysis of different loops related genes, the y-axis represent the number of genes in each GO term. (*D*) Genome browser snapshot of chromatin loop with the annotation of the end anchors at *NCEH1, MB21D2* and *NXPE3* locus, the brown bar represent the enhancer domain of gene, green bar represent the promoter domain and the blue bar is the distal enhancers.

**Figure S5.**
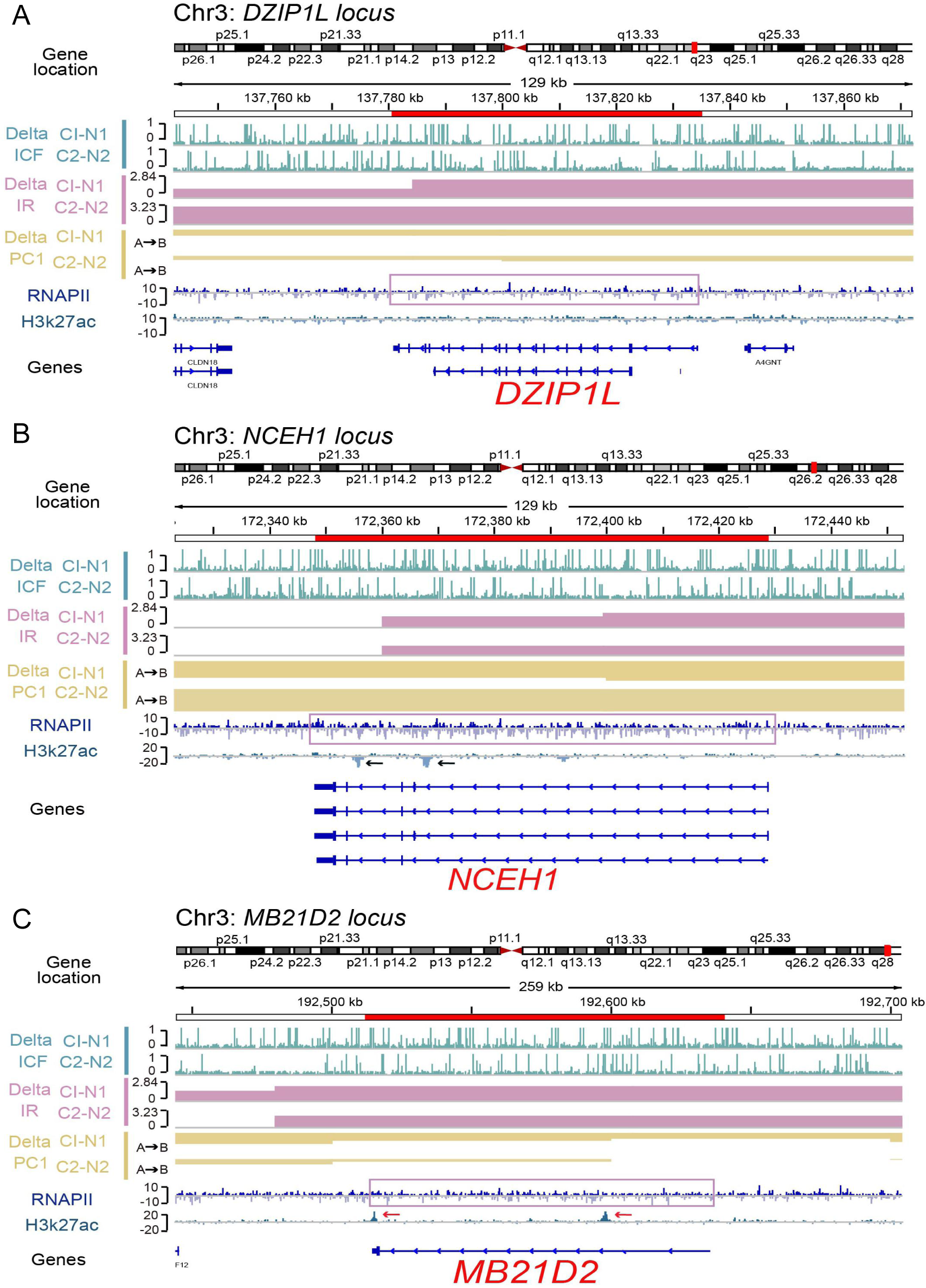
5 most important genes in advanced structure variations in LUAD. Genome browser tracks of ChIP-seq (RNAPII/H3K27ac), compartments changes (Delta PC1), TAD changes (Delta IR) and loop changes (Delta ICF) in tumor compared with tumor tissue at the *NCEH1* (*A*), *DZIP1L* (*B*) and *MB21D2* (*C*) genes locus.

## Notes

### Competing Interest Statement

The authors have declared no competing interest.

